# Metagenomic prediction of methane emissions in sheep using single- and multi-matrix BLUP models with taxonomic and functional microbial features

**DOI:** 10.64898/2026.06.10.731298

**Authors:** Yuhao Li, Chian Teng Ong, Seema Yadav, Michael Aldridge, Peter Fitzgerald, Julius van der Werf, Loan To Nguyen, Elizbeth M Ross

## Abstract

**Background:** Enteric methane emissions from ruminant livestock represent a major greenhouse gas contributor, yet identification of high- and low-emitting ruminants remains expensive and logistically challenging for agricultural methane mitigation strategies. Ruminal microbial profiles derived from long-read sequencing technology provide a potential proxy to predict methane production. The optimal bioinformatic pipelines for processing long-read metagenomic data to perform methane predictions have yet to be determined. Here we evaluated how different metagenomic analysis pipelines affect methane predictive model accuracy in grazing sheep.

**Results:** We applied three bioinformatic pipelines to characterize the taxonomic and functional features of rumen microbiomes from 396 sheep. Functional abundance features were annotated from Clusters of Orthologous Genes (COG) or Kyoto Encyclopedia of Genes and Genomes (KEGG) pathways. The single-matrix model using COG features achieved the highest microbiability (*m*^2^ = 0.942: proportion of variance component explained by microbial features) and predictive accuracy (5-fold cross validation *r* = 0.609: Pearson’s correlation between predicted and observed values). Both functional features outperformed all taxonomic features across all three pipelines in predictive accuracy. The multi-matrix models combined functional and taxonomic features slightly improved methane predictive accuracy across both 5-fold cross-validation and leave-one-day-out validation compared to the models using functional features alone.

**Conclusions:** These findings demonstrate the potential advantages of using long-read metagenomic data to predict enteric methane emissions in ruminants. COG-based functional features achieved the highest predictive accuracy among all feature types, suggesting that functional annotation of existing long-read sequences is sufficient for accurate methane prediction without requiring complementary taxonomic data.

## Background

Enteric methane emissions from ruminant livestock are a significant anthropogenic source of greenhouse gas, contributing 16.8% to 22% of global greenhouse gas emissions (Lean & Moate, 2021). Enteric methane originates as a metabolic byproduct of cellulose digestion and fermentation during ruminant digestion. Rumen methane production is influenced by multiple factors, such as animal physiology (Hickey et al., 2022), dietary composition (Rossi et al., 2022), feeding patterns (Sahu & Arya, 2024), environmental conditions (Palangi et al., 2022), and host genetic characteristics and ruminal microbiome diversity (Difford et al., 2018). While dietary adjustment and nutritional management can reduce ruminant methane emissions in the short term (Eugène et al., 2021), selective breeding for low methane production animals through genomic selection provides potential long-term methane mitigation (Lassen & Difford, 2020). However, phenotyping enteric methane emissions in large numbers of grazing animals, required for accurate genomic selection, is highly challenging.

One potential solution to the challenge of phenotyping large numbers of animals for methane emissions is biomarker-based proxy traits. Effective predictive proxies must be robust and consistent in the detection of high and low methane emission animals. Metagenomic prediction of enteric methane emissions, which uses sequencing data from the rumen microbiome to predict enteric methane emissions, have potential as a proxy trait (Difford et al., 2018; Ross et al., 2020; Ross et al., 2013). Microbiome-based methane prediction has been explored in both cattle and sheep using various microbial feature characterization approaches, including taxonomic and functional profiles (Alemu et al., 2025; Bilton et al., 2025; Martínez-Álvaro et al., 2022; Ross & Hayes, 2022; Sepulveda et al., 2025). Although recent advances in metagenomic research suggest that the functional annotation profiles more directly link to methane production phenotypes compared to using taxonomy profiles (Andersen et al., 2021), the effectiveness of different microbial taxonomic and functional abundance information in predicting ruminal methane phenotypes has not been thoroughly investigated.

Long-read sequencing has advanced the precision of taxonomic and functional quantification (Lam & Ye, 2019; Marić et al., 2024; Mi et al., 2024), but its effect on metagenomic predictions is unknown, as most metagenomic prediction studies have used short read technology. The longer read length and complex genomic region accessibility of long-read data make it valuable for providing high-resolution taxonomic profiles, sensitively quantifying microorganisms and identifying associated functional regions (Portik et al., 2022). This added precision may improve the reliability of metagenomic prediction models by better capturing microbial diversity. Currently, various bioinformatic pipelines relying on different databases are available for identification and annotating long-read metagenomic data, including direct read-based classification and various functional mapping workflows (Kim et al., 2024). However, these approaches have not been compared in their ability to predict methane emission phenotypes across diverse cohorts.

Here we used long-read sequence data from ruminal fluid of 396 sheep to test the effectiveness of different bioinformatic approaches for metagenomic predictions. Three different bioinformatic approaches were used to generate quantitative microbial taxonomic and functional abundance matrices for calculation of microbiability and prediction accuracy. We also tested the effect of integrating both microbial taxonomic and functional matrices into a multi-matrix methane prediction model.

## Methods

All animal experiments and measurement approaches were approved by the University of New England’s Animal and Ethics Committee under authority number: ARA-2024-1549-2323 / ARA23-013.

### Experimental Protocol

504 lambs from the 2022 drop Meat & Livestock Australia (MLA) Resource Flock at Kirby in Armidale, New South Wales, Australia were measured. Animals were managed under free-grazing conditions throughout the study period. All sheep were fasted one hour before each measurement to minimize short-term dietary effects. Enteric methane production measurements were recorded over 7 days in March 2023. A subset of 396 animals, including 188 female and 208 males between 5.48 and 6.06 months of age (mean ± standard deviation: 5.75 ± 0.12 months, median: 5.75 months), with both methane emission data and rumen fluid samples were used in the analysis. On average, 72 animals per day were assessed with 12 Portable Accumulation Chambers (PACs) operating simultaneously. The similar age range of sheep was selected to minimize individual differences in rumen development stages and maintain consistent metabolic states. Rumen samples were collected on all animals immediately post PAC measurement and a subset of total 396 animals with both methane emission data and rumen fluid samples were used in the subsequent analysis.

### Methane Emission Measurement Protocol

Methane production (mL/min) was measured over seven consecutive days using PACs. Following established PAC methodology protocols, individual animals were weighed immediately before chamber entry using precision scales. Each animal was placed in a PAC for a 50-minute measurement period. Methane production was recorded in milliliters per minute (mL/min). All methane production was quantified at the terminal time point using Eagle2 infrared spectroscopy (RKI Instruments, USA), a methane-specific infrared sensor (0-100% volume with auto-ranging capability to ppm). All measurements were conducted under standardized reference conditions. Between each measurement, chambers were “refreshed” using a garden blower and a 30-minute chamber equilibration period was implemented. These 30-minute intervals ensured atmospheric conditions within PAC returned to baseline environmental levels before the next animal entry. Chamber allocation and temporal variables were systematically recorded. Data included methane animal identification numbers, body weight (kg), sex, individual chamber numbers, date of measurement and precise entry/exit timestamps for each measurement. Background atmospheric methane production (mL/min) was measured before and after each session. Corresponding environmental parameters including temperature (°C), humidity (%), and atmospheric pressure (kPa) were recorded simultaneously at the 50-minute measurement point. After completion of each 50-minute methane measurement, animals were rumen sampled using an oral stomach tube and catheter syringe. Post rumen sampling, animals were returned to their paddock to resume normal grazing behavior, ensuring minimal disruption to sheep management practices.

Animals entered chambers in randomized sequences across six daily runs. Each methane measurement run consisted of a batch of 12 animals measured simultaneously, designated as T_0_ to T_5_. PAC entry times were T_0_ (08:00-08:16), T_1_ (09:30-09:46), T_2_ (11:00-11:16), T_3_ (12:30-12:46), T_4_ (14:00-14:16), and T_5_ (15:30–15:46). Approximately 1.5-hour time intervals separated each methane measurement run. Each run allowed a 16-minute window for animal placement. Each run was assigned a time interval value, calculated as hours from the daily baseline (08:00) to each animal’s chamber entry time. Spearman’s rank correlations were used to assess the relationship between run and methane production within each measurement day. As methane production showed mild departure from normality (Shapiro-Wilk W = 0.983, P < 0.001), consistency of temporal trends across days was assessed using aligned rank transform analysis of variance (ART-ANOVA) (Wobbrock et al., 2011) to test day × run interaction effects. All statistical analyses were conducted in R version 4.4.0 (R Core Team, 2024).

### Environmental Correction of Methane Production

Environmental conditions varied daily, requiring methane production corrections for accurate methane emission comparisons. Methane production was corrected for daily air temperature (*T*_air_, °C) and atmospheric pressure (*P*_air_, kPa) and relative humidity (*RH*_*air*_, %) using environmental factors. Firstly, the dry air pressure (*P*_dry_, kPa) was calculated from saturated vapor pressure following Magnus-Tetens formula (Alduchov & Eskridge, 1996) and corresponding to relative humidity calculation by

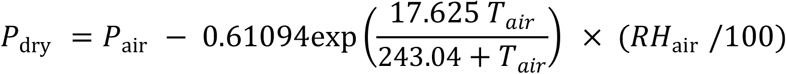

where *T*_air_, *P*_air_, *RH*_air_ were measured simultaneously with each methane production measurement. Next, net methane production was calculated by subtracting background methane production (*C*_bg_) from measured production (*C*_air_), and corrected methane production (*C*_STP_) standardizing by standard temperature and pressure conditions (STP) according to the ideal gas law (McNaught & Wilkinson, 1997) :

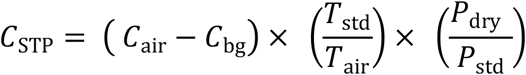

where *T*_std_ was the standard temperature (273.15 K), *T*_air_ was the measured temperature in Kelvin (measured °C + 273.15), *P*_dry_ was dry air pressure (kPa), *P*_std_ was the standard atmospheric pressure (101.325 kPa). This formula was used as a available correction factor to the methane production rate (*ml*/*min*) measured by the PAC approach, ensuring the methane phenotype was standardized to STP conditions for all subsequent analyses.

### Sample collection and DNA extraction

Immediately following methane measurement in the PACs, the sheep were moved to the sampling station, where rumen fluid was obtained from all individual animals. Approximately 20 mL of fluid was extracted using an oral stomach tube and transferred into two 10ml tubes and immediately stored in a freezer until processing. In total, 396 rumen fluid samples were used in this study.

Metagenomic DNA from each rumen sample was extracted using QIAamp PowerFecal Pro DNA Kit (QIAGEN, Germany). The DNA samples were shipped to the University of Queensland at 4°C for downstream analyses. Sequencing library was prepared using the Native Barcoding Kit 96 V14 SQK-NBD114.96 (Oxford Nanopore Technologies, United Kingdom), following the manufacturer’s protocol. Briefly, the extracted DNA was added to the end-repair reaction mix and subsequently ligated with a unique native barcode (between NB01-NB96) in each sequencing run. To address barcode imbalance and optimize sequencing performance, the 396 rumen microbiome samples were pooled in equimolar ratios and sequenced across 10 separate runs, with 38–40 samples per run. This approach maximized flow cell usage while ensuring the required minimum of 1 million reads per sample. Each of the pooled libraries was quantified using Qubit dsDNA HS Assay Kit (ThermoFisher Scientific, USA) and loaded on a FLO-PRO114M flow cell with R10.4.1 chemistry prior to sequencing with PromethION 2 Solo (Oxford Nanopore Technologies, UK).

### Long-read sequencing and basecalling

Basecalling was performed using Dorado v0.7.0 (Oxford Nanopore, 2023) with the “Super Accuracy” model (v4.3.0). After trimming off the adapters, the remaining reads were demultiplexed by sample barcode and quality-filtered using NanoPack2 (De Coster & Rademakers, 2023) to select for reads with read length >200 bp and Phred score >10. Host DNA contamination was removed by aligning reads to the *Ovis aries* (ARS-UI_Ramb_v3.0) reference genome using minimap2 (Li, 2021). Non-host reads were retained for metagenomic analysis.

### Taxonomic classification and functional annotation with SqueezeMeta

Non-host long-read metagenomic data were taxonomically classified and functionally annotated using SqueezeMeta v1.6.3 (Tamames & Puente-Sánchez, 2018). The automated ‘sqm_longreads.pl’ pipeline of SqueezeMeta was used to process 396 clean FASTQ long-read data files. This pipeline utilized Prodigal v2.6.3 (Hyatt et al., 2010) to predict open reading frames (ORFs) for individual reads, with translated protein sequences obtained for protein alignment. Protein alignment was processed by using DIAMOND v2.1 (Buchfink et al., 2021) against the NCBI GenBank non-redundant protein database (NCBI-NR). Taxonomic classification was determined using the lowest common ancestor algorithm to assign aligned individual reads from phylum to genus levels. The genus-level classification threshold of 85% protein similarity (default) was used. Only taxa belonging to *bacteria, archaea*, and *eukaryote* kingdoms were retained for subsequent analysis. Functional annotation of metagenomes employed homology alignment results against the Kyoto Encyclopedia of Genes and Genomes database (KEGG version 110) (Kanehisa & Goto, 2000) and eggNOG v5.0 database (Huerta-Cepas et al., 2019), which contains clusters of orthologous groups (COG) functional classifications.

In total, seven abundance matrices were constructed from the SqueezeMeta pipeline: five representing taxonomic levels (phylum to genus) and two representing functional categories: genes and genomes functional pathways (KEGG) and Clusters of Orthologous Groups of proteins (COG). These matrices provide a quantitative representation of microbial community taxonomic diversity and functional annotation patterns.

### Taxonomic classification with Kraken2

Taxonomic classification with Kraken2 is performed using direct DNA-DNA by aligning k-mers against microbial reference databases. Therefore Kraken2 (Wood et al., 2019) was used as a taxonomic classification approach that differs essentially from the SqueezeMeta pipeline, which uses a DNA-to-protein alignment strategy. Additionally, since the SqueezeMeta pipeline is relatively slow and resource-intensive, Kraken2 was evaluated as an alternative with faster processing speed and lower resource requirements for taxonomic classification. Two prebuilt Kraken2 format databases were employed: NCBI nucleotide database (NCBI-NT) and the Genome Taxonomy Database (GTDB) v226 (Parks et al., 2021) for high-resolution *prokaryotic* classification (*bacteria* and *archaea*). To optimize sensitivity for long-read sequencing data and enable detection of low-abundance taxa and potential contaminants, the confidence threshold was customized to 0.005 in Kraken2 (Marić et al., 2024). This customization ensured an appropriate balance between sensitivity and sufficient specificity for reliable Kraken2 taxonomic classification. This process generated taxonomic abundance matrices for Kraken2-NT and Kraken2-GTDB methods at 5 taxonomic ranks (phylum, class, order, family, genus).

### Methane phenotype and covariates

Methane production (*ml*/*min*) was used as the response variable across all prediction models. Linear regression was used to assess the association between enteric methane production and multiple variables including body weight, sex, age, chamber and time of run. Body weight and time of run were identified as covariates by significant level *p* -value less than 0.05.

### Normalization of microbial abundance matrices and variables standardization

Low-abundance microbial taxonomic and functional features (total count < 10 reads) were removed to reduce noise. To account for uneven sequencing depth, all microbial feature matrices were normalized to relative abundances, including taxonomic profiles (five levels from both SqueezeMeta and Kraken2) and functional profiles (KEGG and COG from SqueezeMeta). Microbial feature counts were converted into percentages within each sample, with all features summing to 100%. This approach accounted for differences in sequencing depth while preserving the proportional relationships among taxa and functional categories.

Corrected methane production, selected covariates (body weight and time of run), and normalized microbial feature abundance matrices were standardised using Z-score prior to statistical analysis and model fitting.

### Principal Coordinate Analysis and Correlation with Methane Production

Principal coordinate analysis (PCoA) was used to investigate the correlations between microbial abundance features and methane production. Microbial Bray-Curtis distance matrices were calculated based on genus-level taxonomic abundance and from KEGG- and COG-level functional profiles. PCoA was performed on each distance matrix. To capture the major gradients in microbial feature composition, the first five principal coordinates were used to estimate Spearman’s rank correlation coefficients with residual methane production. Methane production residual values were calculated by using body weight and time of run in linear regression. Statistical significance was defined as *p* < 0.05.

### Single-matrix BLUP models

Best linear unbiased prediction (BLUP) models were implemented using ASReml-R 4.2 (Butler & Cullis, 2007) in R 4.4.0 to evaluate the predictive value of microbial features for variation in methane emissions, following the framework of Ross et al. (2013). In first instance, we fitted one feature as a random effect, referred to as single matrix model.

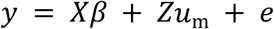

Where *y* is the vector of methane production,*β* is the vector of fixed effects, *Z* is the incidence matrix linking samples to microbial random effects, *u*_*m*_ is the vector of random microbial effects with 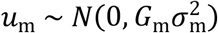, and *e* is the vector of residuals with 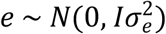, and I is the identity matrix, 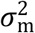 is represents the microbial variance component (calculated from taxonomic or functional features), and 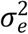 is the residual variance. The microbial relationship matrix *G*_*m*_ was calculated as:

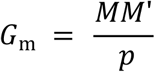

Where *M* is the standardized microbial feature matrix (*n* samples × *p* features), *M*’ is the transpose of *M*, and *p* is the number of microbial features (Additional file 1: Table S1 for the number of microbial features at each taxonomic and functional level). The microbiability *m*^2^ indicating the proportion of methane variance explained by microbial feature effects was calculated as

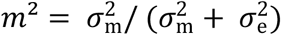

Delta approximation method was employed to calculate standard errors of microbiability estimates (Yadav et al., 2021).

Each microbial data source was evaluated independently using single-matrix BLUP models: five taxonomic levels (phylum to genus) from each of the three pipelines (SqueezeMeta, Kraken2-NT, Kraken2-GTDB) and two functional matrices (COG and KEGG) from the SqueezeMeta pipeline, resulting in total 17 models. In each model, the microbial relationship matrix was fitted as a random effect. Mantel tests based on Spearman rank correlations were used to assess the similarity among all 17 microbial relationship matrices.

### Multi-matrix BLUP models

Multi-matrix BLUP models were used to jointly incorporate taxonomic and functional microbial abundances by fitting multiple random effect components. This modeling approach increases the ability to capture microbial information when taxonomic and functional features explain different and complementary parts of the phenotypic variance. The multi-matrix BLUP models were defined as:

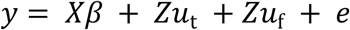

where y is a vector of methane production, *X* and *β* represent the same fixed effects using in the single matrix models, *Z* is the incidence matrix linking samples to random effects. The vectors *u*_t_ and *u*_f_ represent the taxonomic and functional microbial random effects, respectively. The taxonomic and functional random effects were assumed as: 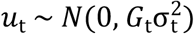 and 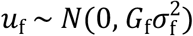. The taxonomic microbial relationship matrix (*G*_t_) was constructed as:

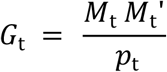

where *M*_t_ is the standardized taxonomic abundance matrix (n samples × *p*_t_ features) at the taxonomic level derived from the SqueezeMeta and Kraken2 pipelines. Similarly, the functional microbial relationship matrix. *G*_f_ was defined as:

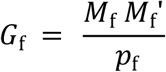

where *M*_f_ is the standardized functional abundance matrix (n samples × *p*_f_ features) derived from KEGG or COG categories. Variance components in the multi-matrix BLUP models decomposed total phenotypic variance into the independent components corresponding to taxonomic variance 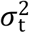, functional variance 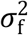, and residual variance 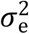. These components 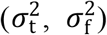 quantify the unique contribution of each microbial feature set to variation in methane emissions, while 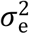 represents the residual variance. Total microbiability (*m*^2^) was calculated as: 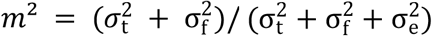.

### Cross-Validation for model evaluation

To evaluate predictive performance, two cross-validation strategies were implemented to assess the accuracy of methane prediction models: random 5-fold cross-validation and leave-one-day-out cross-validation. For random 5-fold cross-validation, animals were randomly divided into five folds with approximately 79-80 animals per test set and 316-317 animals per training set. Each fold served as the test set once, with the remaining four folds used for model training. Consistent partitions across all model types were used to ensure comparability. In addition to the 5-fold cross-validation, leave-one-day-out was also used to assess prediction accuracy. For the leave-one-day-out strategy, each cross-validation iteration excluded all samples on a single day (51-62 animals) for testing, while the remaining six days (334-345 animals) were used for model training.

For each cross-validation fold, all continuous variables (corrected methane production, weight, and run) were standardised using the mean and standard deviation (SD) calculated exclusively from the training set. The calculated standardization parameters were applied to the corresponding test data. Model fitting was performed on the training set phenotypes. In both 5-fold and leave-one-day-out validation, all preprocessing and model training were restricted to the training folds.

Predictive performance was evaluated using the Pearson correlation coefficient (*r*) between predicted and observed standardised methane production, with root mean square error (RMSE) reported as an additional error metric. Mean accuracy (*r*) and RMSE were computed across folds, and fold-specific values were also retained. Standard errors were calculated from the fold-to-fold variation as the standard deviation divided by the square root of the number of folds (*k*), where *k* =5 for 5-fold CV and *k* = number of days for leave-one-day-out CV. Visualization was performed using ggplot2 (Wickham, 2011). All prediction accuracy results are reported as mean ± standard error. Feature-wise random shuffling of all microbial abundancy matrices was performed as a negative control (see Additional file 3: Supplementary Methods for details).

## Results

### Methane production phenotype regression analysis

The mean (± SD) methane production across the 396 sheep were 7.52 ± 2.82 mL/min, ranging from 1.44 to 16.87 mL/min (Fig. 1). The median methane production was 7.20 mL/min, indicating a slightly right-skewed distribution.

**Fig. 1.**
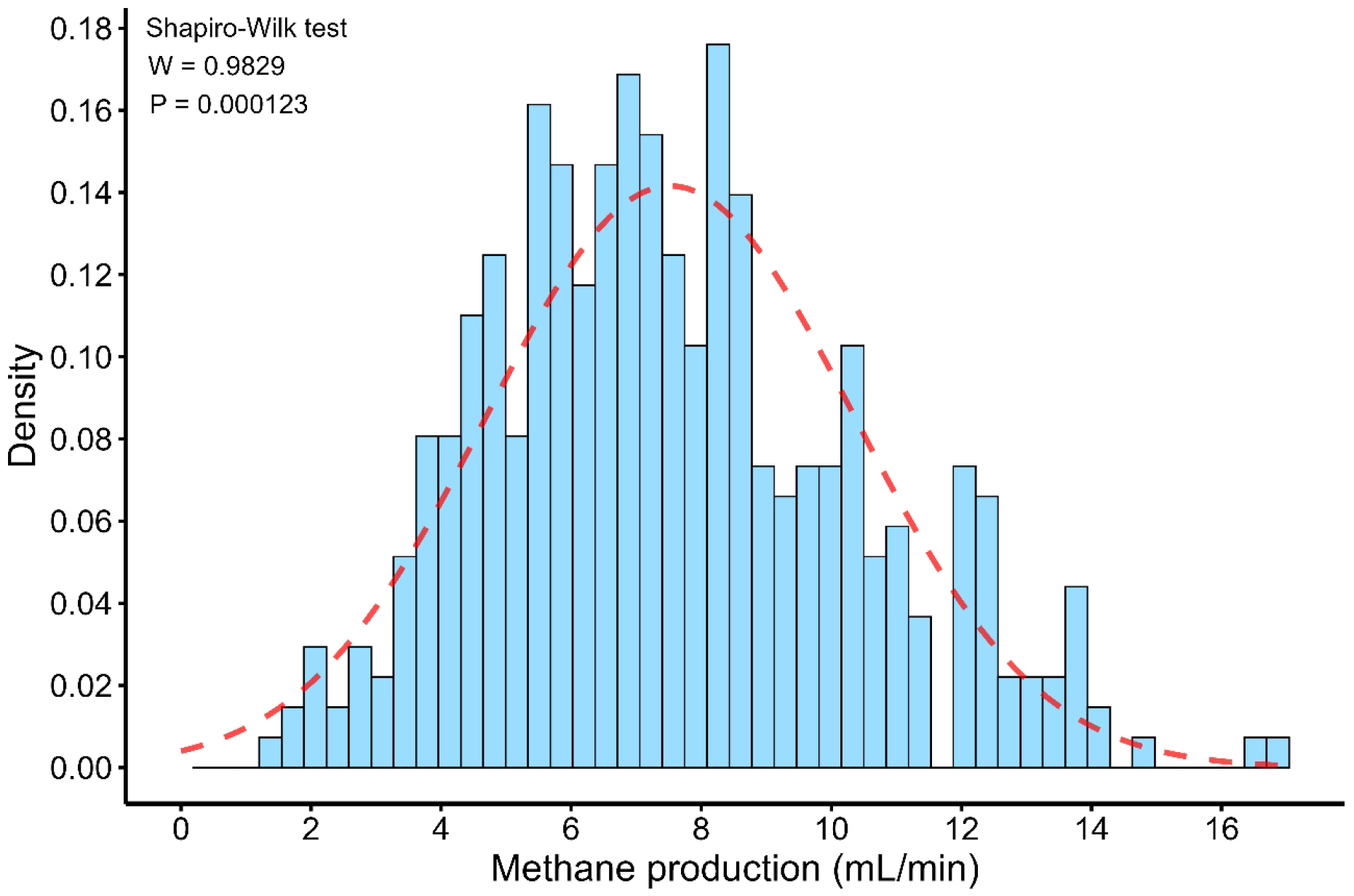
Distribution of methane production sheep measured using portable accumulation chamber (PAC). Methane production (mL/min) was recorded over a 7-day period from 396 animals. The histogram presented the probability density for data normality assessment and showed y-axis with the individual means with fitted normal distribution curve (red dashed line).

Linear regression models (Table 1) showed sex, age and chamber effects had no significant effect on methane production (*R*^2^ < 0.01, *P* > 0.1). Body weight showed a positive association with methane production, accounting for 14.92% of variance, while measurement run explained 35.80% of variance (both *P* < 0.001, Table 1). Together body weight and run explained 52.74% of total variance (*P* < 0.001), confirming their inclusion as fixed effects in all subsequent methane prediction modelling.

**Table 1.** Linear regression analysis of fixed effects on methane emissions in sheep. The proportion of variance explained (*R*^2^) and statistical significance (*p*-value) are shown. *** p < 0.001, ns p > 0.05.

| Factor | $R^2$ | $p$-value |
| --- | --- | --- |
| Body Weight (kg) | 0.1492 | <1.53e-15 (***) |
| Sex | <0.0001 | 0.912 (ns) |
| Age (months) | 0.0006 | 0.625 (ns) |
| Chamber | 0.0036 | 0.232 (ns) |
| Time of run (hour) | 0.3580 | <2e-16 (***) |
| Body Weight (kg) + Time of run (hour) | 0.5274 | <2e-16 (***) |

Methane production consistently decreased from T_0_ to T_5_ within each experimental day, and this pattern was observed across all sampling days. The raincloud plots (**PCoA**Fig. S1) showed that methane production peaked at T_0_ (median: 8.2-13.1 mL/min) and reached their minimum at T_5_ (median: 3.4-9.7 mL/min). ART-ANOVA revealed that both sampling day (*F*_[6,354]_ = 20.55; *p* < 2×10^−16^) and run (*F*_[5,354]_= 58.75; *p* < 2×10^−16^) had significant effects on methane production. The day × run interaction (*F*_[30,354]_ = 1.58; *p* > 0.03) was not significant, indicating that temporal decline patterns were consistent across all 7 days. Spearman’s rank correlations revealed consistent negative relationships between run and methane production within each day (*ρ*= -0.422 to -0.758, all *p* < 0.001), showing a repeatable decreasing trend across all sampling days.

### Long-read metagenomic sequence data

Long-read metagenomic data profiles from sheep rumen samples were described using descriptive statistics (mean ± SD). On average 27.3% (± 5.7%) of raw reads were discarded as they failed to meet minimum quality thresholds (Phred score >10 and read length > 200 bp). The resulting high-quality dataset comprised 1.68×10^6^ (± 1.08×10^6^) reads per sample, with an average read length of 643 (± 222) bp. The average N50 read length was 1,014 (± 1,013) bp, indicating substantial variation in read length distribution across the 396 samples.

We compared the model performance of three metagenomic pipelines (SqueezeMeta, Kraken2-NT, and Kraken2-GTDB) for methane prediction. Among the three methods in taxonomic classification, Kraken2-GTDB consistently detected the most taxonomic features, with 30192 genera detected. Principal coordinate analysis (PCoA) clustering results (Fig. 2A, Additional file 2: Fig.S2 and Additional file 1: Table S1) showed that for the COG features PCoA1 and PCoA2 accounted for 59.18% and 6.2% of the variance, respectively. Linear regression models were fitted with body weight and time of run as covariates. None of the principal coordinates showed significant correlation with methane production after FDR correction (Additional file 1: Table S1).

**Fig. 2.**
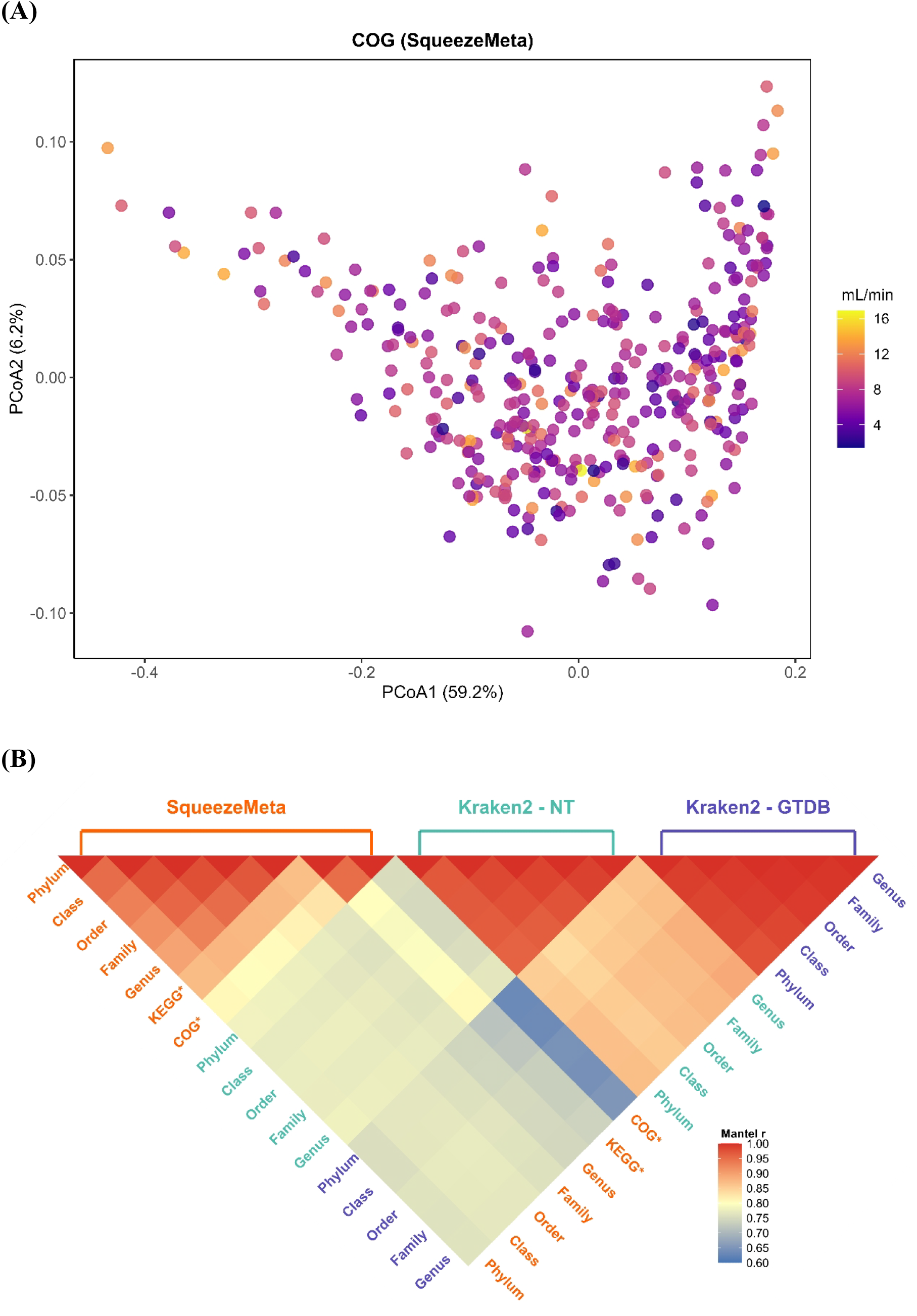
**(A) Principal coordinate analysis (PCoA) of rumen microbial COG abundance of SqueezeMeta associated with methane production phenotypes**. Colorful points represent individual samples colored by methane production (mL/min), showing the distribution of microbial features across the methane production gradient. Color scale ranges from purple (lowest emitters, ∼ 4 mL/min) through pink (intermediate emitters) to orange (highest emitters, ∼16 mL/min). PCo1 and PCo2 explain 57.0% and 14.8% of the total variance, respectively. **(B). Mantel test correlations (***r***) between microbial relationship matrices**. Heatmap of pairwise Mantel correlation coefficients (*r*), calculated between microbial relationship matrices constructed from five taxonomic levels (phylum, class, order, family, genus) and two functional annotations (KEGG* pathways, COG* categories) across three metagenomic approaches: SqueezeMeta (orange), Kraken2-NT (green), and Kraken2-GTDB (blue). Mantel *r* value (approaching 1) indicates higher concordance in microbial feature structure as captured by the corresponding abundance matrices.

Pairwise Mantel tests were used to evaluate the similarity of microbial relationship matrices generated from different taxonomic and functional abundance profiles. Overall, similar structures were observed across all microbial relationship matrices, with strong positive Spearman’s rank correlation coefficients Mantel *r* ranging from 0.63 to 0.99 (all *P* < 0.0001; Fig. 2B). Functional microbial relationship matrices (KEGG and COG) showed the lowest correlation with all microbial taxonomic relationship matrices (Mantel *r* < 0.8; Fig. 2B). The COG-based matrix showed the lowest correlation (Mantel *r* = 0.63 - 0.65) to taxonomic microbial relationship matrices from the Kraken2-GTDB approach (Fig. 2B). Microbial relationship matrices generated using the SqueezeMeta pipeline (COG vs genus, Mantel *r* = 0.83; KEGG vs genus, Mantel *r* = 0.87; Fig. 2B) correlated to each other more strongly than those from Kraken2 pipeline. The taxonomic microbial relationship matrices generated using Kraken2 via different databases (NCBI and GTDB) were very highly correlated (Mantel *r* > 0.8; Fig. 2B).

### Microbiability estimates of single-matrix BLUP models

The contribution of rumen microbial variation to methane phenotype was quantified using microbiability (*m*^2^) with standard error for each approach separately (Fig. 3 and Additional file 1: Table S2). In single-matrix BLUP models, the functional features (KEGG and COG) showed higher microbiability estimates than single taxonomic features across all three pipelines. The microbiability of all single taxonomic matrix models varied by both taxonomic levels and classification approaches.

**Fig. 3.**
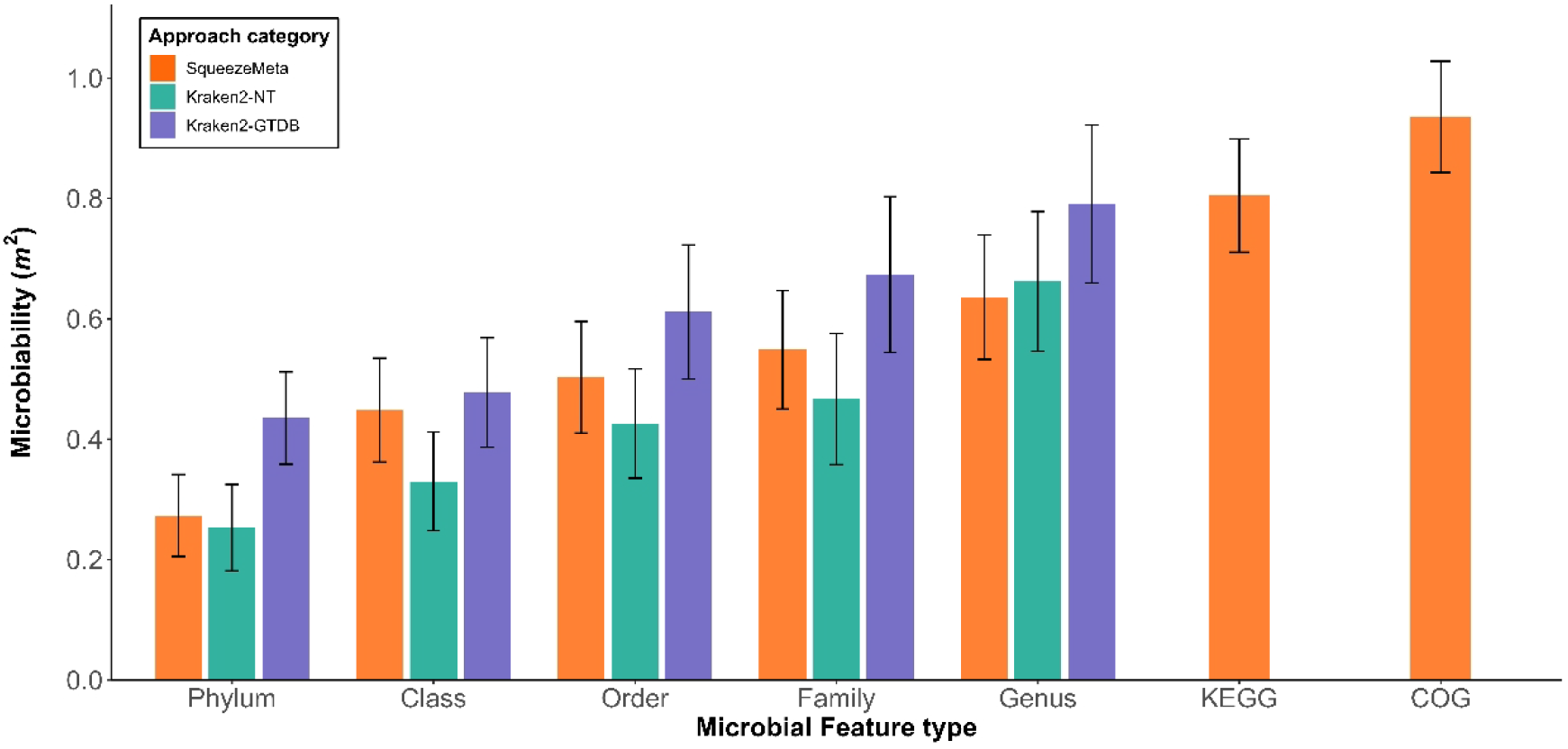
Microbiability estimates from single-matrix BLUP models. Estimates were derived from taxonomic levels (phylum to genus) and functional annotation (KEGG and COG) using three metagenomic approaches: SqueezeMeta, Kraken2-NT, and Kraken2-GTDB. Error bars represented standard errors of microbiabilities.

Functional microbial features, available only from the SqueezeMeta pipeline, demonstrated higher microbiability estimates than taxonomic classifications generated by any of the three pipelines (Fig. 3 and Additional file 1: Table S2). The COG-based microbial relationship matrix had the highest estimated microbiability (*m*^2^ = 0.935 ± 0.092), while the KEGG pathway-based microbial relationship matrix explained less variance (*m*^2^ = 0.805 ± 0.094).

All three taxonomic approaches showed consistent microbiability patterns with values increasing from higher to lower taxonomic levels (phylum to genus), while residual variance components (*V*_e_) decreased correspondingly (Fig. 3 and Additional file 1: Table S2). Models with the genus-level microbial relationship matrix achieved the highest microbiability among all taxonomic levels: SqueezeMeta (*m*^2^ = 0.636 ± 0.104), Kraken2-NT (*m*^2^ = 0.662 ± 0.116), and Kraken2-GTDB (*m*^2^ = 0.791 ± 0.131). Kraken2-GTDB consistently had higher microbiability values than the other two approaches (Fig. 3 and Additional file 1: Table S2). Except for the genus level, Kraken2-NT using NCBI database estimated the lowest microbiability compared with the SqueezeMeta and Kraken2-GTDB approaches. Thus, the significantly higher taxon density (Table 2) from Kraken2-GTDB genus level (*n* = 30192 features) may contribute to increased microbiability.

**Table 2.** Taxonomic and functional microbial features identified by 3 different pipelines.

| Features Identified | Method |  |  |
| --- | --- | --- | --- |
|  | SqueezeMeta | Kraken2-NT | Kraken2-GTDB |
| Phylum | 242 | 185 | 201 |
| Class | 655 | 276 | 615 |
| Order | 1174 | 593 | 2154 |
| Family | 2008 | 1212 | 5925 |
| Genus | 3042 | 3156 | 30192 |
| KEGG | 9591 | - | - |
| COG | 22873 | - | - |

### Multi-matrix BLUP Variance components

We fitted BLUP models with two random effects, both taxonomic and functional microbial relationship matrices to evaluate the importance of feature type by decomposing the variance contributions (Fig. 4 and Additional file 1: Table S3). Functional features (COG or KEGG) accounted for most of the total explained variance in these methane prediction models.

**Fig. 4.**
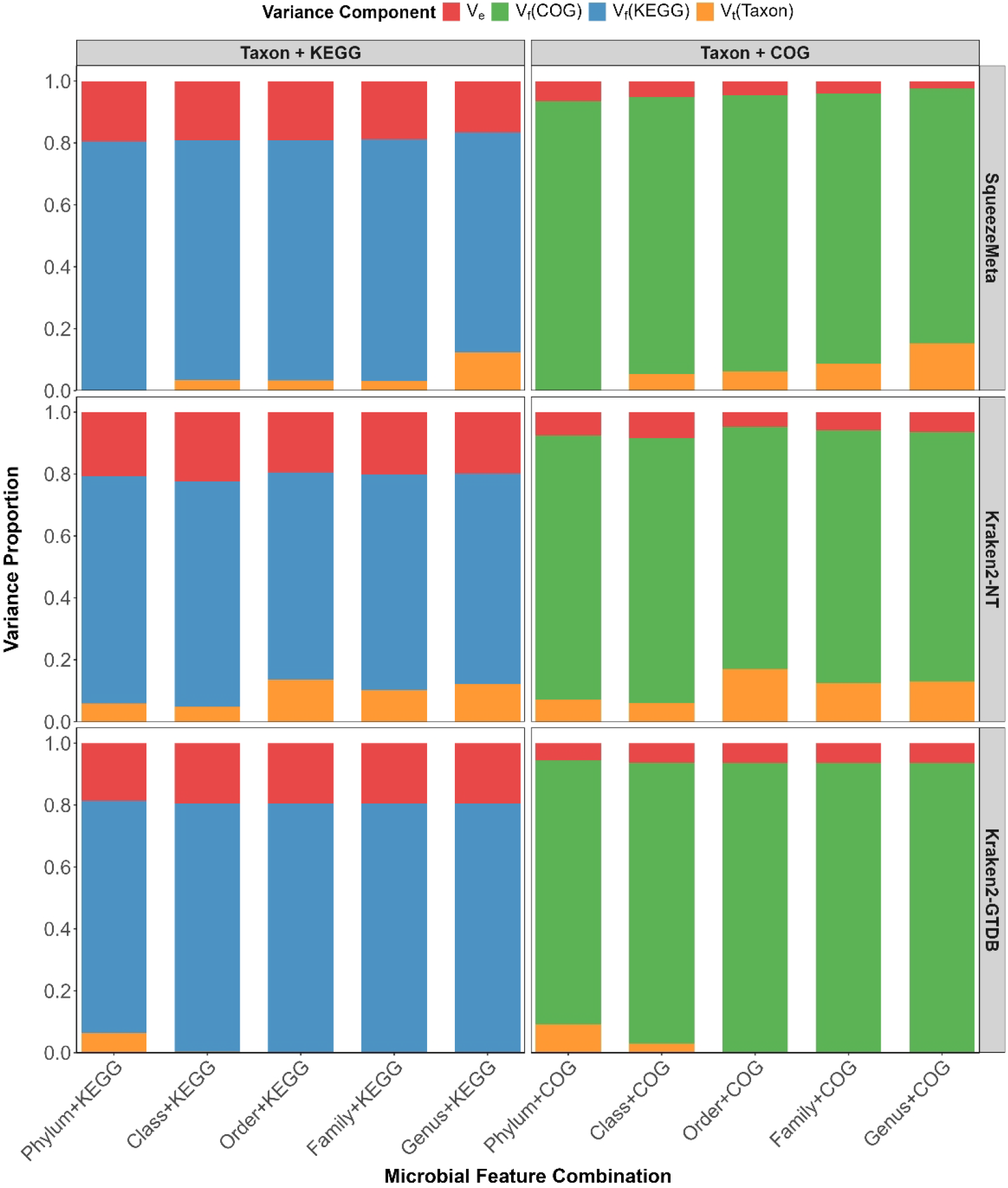
Multi-matrix BLUP models combining taxonomic level (phylum to genus) with functional level (KEGG and COG) refers to additive microbial taxonomy-by-function interaction models. Y-axis shows the proportion of total variance explained by each component contributed to microbiability estimates (Additional file 1: Table S3). The colors indicate variance component categories: *V*_*e*_ (residual variance, 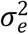), *V*_f_ - COG and *V*_f_ - KEGG (functional variance, 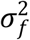), *V*_t_ - Taxon (taxonomic variance,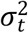).

The microbiability (*m*^2^) estimates from multi-matrix models were higher than those from single-matrix models. The genus + COG models reached the highest total microbiability, *m*^2^ = 0.975 ± 0.086 (SqueezeMeta genus + COG) and *m*^2^ = 0.936± 0.092 (Kraken2-NCBI genus + COG). In the genus + COG model, the functional component (COG) dominated the proportion of total variance explained (82.2%), with *V*_f_ = 0.430 ± 0.071 substantially exceeding the taxonomic component (*V*_t_ = 0.080 ± 0.050), while residual variance remained negligible (*V*_e_ = 0.013 ± 0.045) (Additional file 1: Table S3). Taxonomic and functional microbial relationship matrices were highly correlated (Mantel *r* = 0.87), but total microbiability still showed an increasing trend with the integration of complementary microbial information (*m*^2^ = 0.975 ± 0.086).

The genus + KEGG models reached *m*^2^ = 0.833 ± 0.094 (SqueezeMeta genus + KEGG); *m*^2^ = 0.805 ± 0.094 (Kraken2-GTDB genus + KEGG) (Additional file 1: Table S3). These microbiability values were higher than or comparable to corresponding single functional matrix models (KEGG only: *m*^2^ = 0.805 ± 0.094) respectively (Additional file 1: Table S2).

### Cross validation of methane prediction

Functional feature models achieved the highest methane prediction accuracies in 5-fold validation (mean ± standard deviation across 5 folds). The functional single-matrix models from SqueezeMeta approach estimated higher mean accuracies in COG (*r* = 0.634 ± 0.038, RMSE = 0.806 ± 0.031) and KEGG (*r* = 0.624 ± 0.032, RMSE = 0.815 ± 0.026) models than all taxonomic single-matrix models (Fig.5A and Additional file 1: Table S4). The taxonomic single-matrix model reached the highest predictive accuracy in the SqueezeMeta-based genus-level (*r* = 0.575 ± 0.015, RMSE = 0.860 ± 0.025), outperforming all Kraken2-based taxonomic single-matrix models (Fig.5A and Additional file 1: Table S4). The predictive accuracy of taxonomic single-matrix models increased gradually from phylum to genus across all approaches (Fig.5A and Additional file 1: Table S4).

**Fig. 5.**
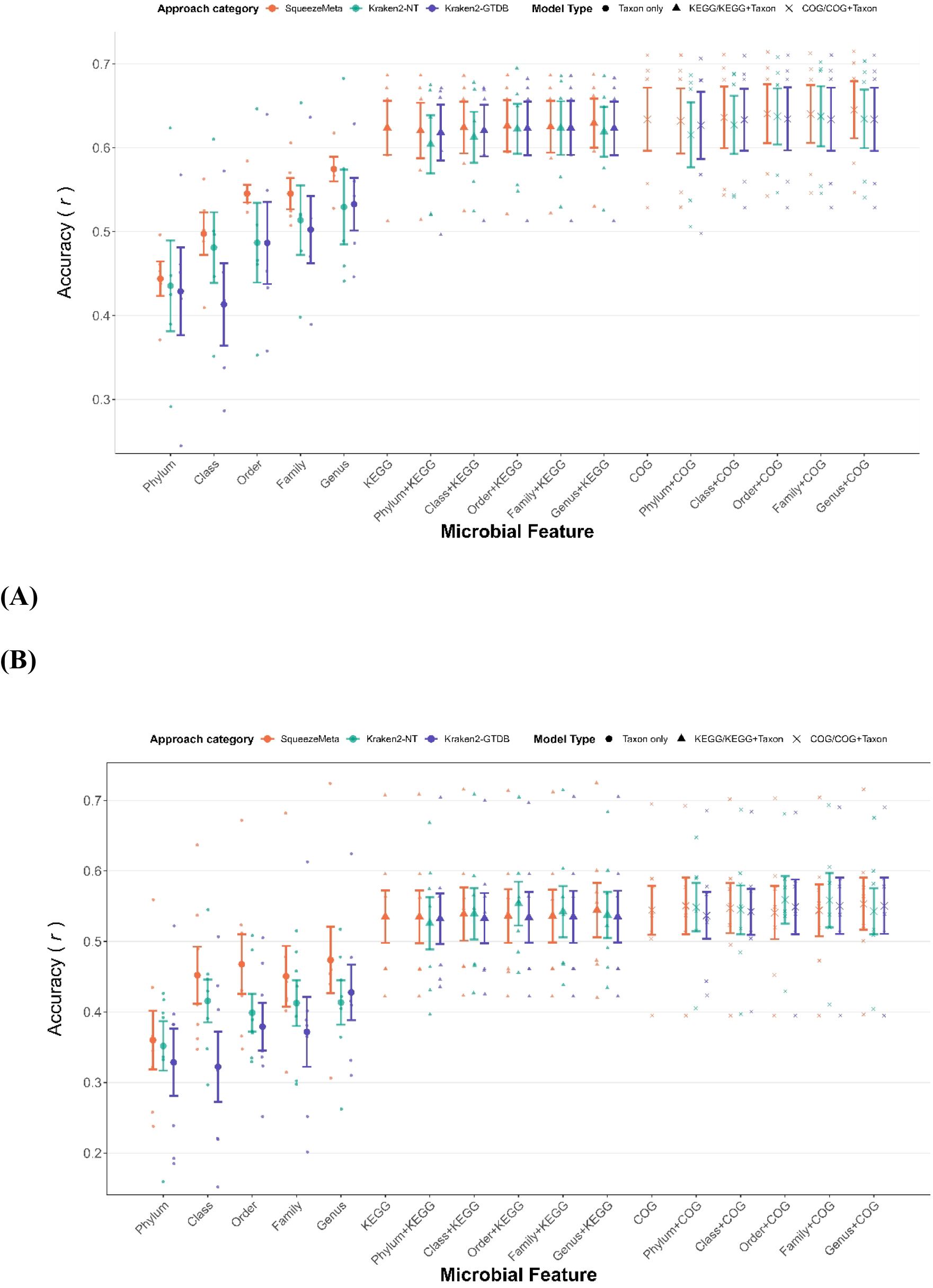
Cross-validation performance of BLUP models for methane prediction. **(A) 5-fold cross-validation; (B) leave-one-day-out validation**. Prediction accuracy is shown as the Pearson correlation coefficient (*r*) between predicted and observed methane production, with error bars (mean ± standard error across validation folds. Single-matrix models (circles) include single taxonomic features, whereas multi-matrix models combined taxonomic features with KEGG (triangles) or COG (crosses) functional annotations. Results are shown for three metagenomic approaches: SqueezeMeta (orange), Kraken2-NT (teal), and Kraken2-GTDB (purple) (n = 396 sheep).

Slight or no improvement over the functional single-matrix models was observed in the multi-matrix predictions (Fig.5A and Additional file 1: Table S4). The highest accuracy multi-matrix model (SqueezeMeta genus + COG) achieved *r* = 0.645 ± 0.034 (RMSE = 0.800 ± 0.030) representing only a marginal increase (Δ *r* = + 0.011, ∼1.7%) compared to the COG single-matrix model (Fig.5A and Additional file 1: Table S4). In contrast, the multi-matrix models Kraken2-NT genus + COG (*r* = 0.634 ± 0.035, RMSE = 0.806 ± 0.029) and Kraken2-GTDB genus + COG (*r* = 0.634 ± 0.038, RMSE = 0.806 ± 0.031) exhibited no mean accuracy improvement compared to their corresponding COG single-matrix model (*r* = 0.634 ± 0.038, RMSE = 0.806 ± 0.031) (Table Additional file 1: S4). Similarly, multi-matrix models with taxonomic features + KEGG combinations showed consistent patterns with taxonomy + COG, which indicated no improvement or even reductions in accuracy (Fig.5A). Accuracy differences between single- and multi-matrix models were consistently small overall from 5-fold random validation. These accuracy comparison results suggested that functional features capture the essential predictive capability for methane phenotypes.

Prediction accuracies of leave-one-day-out validation in all taxonomic models were 9.2% to 25.9% lower than equivalent 5-fold validation models (Fig.5A-B and Additional file 1: Table S4-S5). Functional features demonstrated greater robustness in model performance from 5-fold to leave-one-day-out validation. COG only declining by 14.2% (from *r* =0.634 ± 0.038 to *r* = 0.544 ± 0.035) and KEGG only by 14.3% (from *r* = 0.624 ± 0.032 to *r* = 0.535 ± 0.037) compared to the equivalent 5-fold validation models (Additional file 1: Table S4-S5).

Multi-matrix predictions were subsequently evaluated. The SqueezeMeta genus + COG multi-matrix model achieved *r* = 0.645 ± 0.034 in 5-fold validation, which declined by 14.3% to *r* = 0.553 ± 0.037 in leave-one-day-out validation (Additional file 1: Table S4-S5). All prediction accuracy results demonstrated that the dominant microbiability contribution of functional features (COG and KEGG) contributed more to microbiability and predictive accuracy than all levels of taxonomic features, with this pattern consistent across both validation strategies.

As a negative control, we randomly shuffled both taxonomic and functional microbial abundance matrices with microbiability and prediction accuracy dropping to non-significant levels (Additional file 3: Supplementary Result, Table S6).

## Discussion

### Summary

This study presented metagenomic predictions for methane using microbial taxonomic and functional features from sheep rumen samples in a BLUP model framework. We estimated microbiability and prediction accuracy to explore the contribution of different microbial feature categories by the single- or multi-matrix BLUP modeling. We achieved higher prediction accuracies in a 5-fold cross validation across all models (*r* = 0.413 - 0.645) than validation approaches of other studies, which achieved maximum accuracies of 0.216 to 0.557 in sheep (Alemu et al., 2025; Bilton et al., 2025; Ross et al., 2020) of which all used short read data to generate quantitative metagenomic matrices. Benchmarking of bioinformatic approaches demonstrated that functional features consistently outperformed taxonomic features among all BLUP models. Multi-matrix BLUP models provided minimal or no improvement beyond single-matrix models.

### Functional features dominate methane prediction

Microbial functional features (KEGG and COG) achieved higher prediction accuracies compared to taxonomic features. This finding was observed consistently in two cattle populations reported by Sepulveda et al. (2025), that functional features (KEGG and COG) captured methane-associated variation more effectively than taxonomic features. Functional profiles likely provide a more biologically relevant representation of rumen microbial activity, as they reflect metabolic pathways rather than taxonomic abundance alone (Andersen et al., 2021). Fundamental microbiome studies previously highlighted microbial functional features showing robustly associated with host phenotype than taxonomic features (Iablokov et al., 2020; Kuang et al., 2016).

Integrating taxonomic and functional features in multi-matrix models simultaneously provided minimal improvement or even reductions in prediction accuracy. This may suggest redundant information in taxonomic and functional features integration. One broader microbial features integration study indicated that combining multiple feature data does not improve accuracy once the essential information is captured in the model (Lu et al., 2005). This suggests that the functional features (COG or KEGG) capture the comprehensive information provided by metagenomic data. The taxonomic approaches likely predict methane emissions because they reflect underlying functional variation. This highlights the value of using “shotgun” metagenomic approaches, compared to amplicon-based approaches which are only able to identify taxa but not directly observe functional variation.

### Balancing precision and efficiency in metagenomic identification tool selection

Bioinformatic approaches for generating microbial abundance matrices from raw reads reflect trade-offs between computational efficiency and the precision of microbial feature detection. Here we observed that the selection of bioinformatic pipelines for microbial feature generation is a more important determinant of methane prediction accuracy than feature abundance resolution alone. In this study, Kraken2, which was selected for its fast processing speed and reasonable memory requirements (Irankhah et al., 2024), processed each long-read fastq uncompressed file (averaging 1.0 GB, range: 2.5 - 7.7 GB) with an average of 3.6 minutes (range: 2.5 - 7.7 minutes) in our pipeline. In comparison, SqueezeMeta, which performed protein alignment steps with DIAMOND and requires dramatically longer processing time and more resources (Agustinho et al., 2024), required 10.2 ± 5.9 hours (mean ± SD) per sample in our pipeline to obtain both functional and taxonomic abundance profiles. Rapid read-to-reference taxonomic classification and abundance matrices generation become essential when processing large sample cohorts. Therefore, while SqueezeMeta outperformed Kraken2 in prediction accuracy, it may not be the best approach in a commercial setting. The choice of bioinformatic pipeline should reflect the balance between accuracy requirements and computational resources, especially for large-scale methane prediction applications.

### Inconsistent relationship between microbiability and prediction accuracy

Microbiability did not consistently reflect corresponding methane prediction accuracy in the single-matrix models. For example, the highest microbiability of Kraken2-GTDB genus single-matrix model did not reach greater accuracy than SqueezeMeta genus single-matrix model. While using single KEGG functional feature showed similar microbiability estimates to Kraken2-GTDB genus-level, KEGG single-matrix model demonstrated higher accuracy in methane prediction. Similarly, Hess et al. (2023) evaluated 2 scenarios with microbial relationship matrices in methane prediction models, demonstrating the similar modeling performance relationship: one scenario achieved high microbiability (*m*^2^ > 0.8) but modest prediction accuracy (*r* < 0.4), while another scenario showed significantly different microbiability (*m* ^2^ < 0.2) maintained similar prediction accuracy (*r* < 0.3). Hence, when using microbiability to assess potential prediction accuracy, care should be taken not to over interpret the results. In this study, the high microbiability estimates (*m* ^2^ > 0.90) observed in both single-and multi-matrix models corresponding to lower prediction accuracies (*r* ≈ 0.5 – 0.6) confirm this concern. Variance component estimation of microbiability from BLUP models may be inflated depending on the matrix construction method or properties of the microbial relationship matrix (Bruijning et al., 2023; Saborío-Montero et al., 2021). This suggests that validated strategies of prediction accuracy may provide a more reliable assessment of methane prediction than considering microbiability.

### Validation strategies reveal model robustness

Different validation strategies led to different predictive performance estimates. Random 5-fold cross-validation consistently led to higher prediction accuracies than the corresponding leave-one-day-out validation using the same pipeline and microbiome feature matrix. The random fold assignment does not account for day-to-day variability. Cross-day methane prediction is particularly challenging, as both microbiome community and methane emissions are influenced by temporal effects. The unmeasured temporal effects captured by leave-one-day-out validation are artificially minimized in random cross-validation (Roberts et al., 2017). This is compatible with the temporal variability observed in PAC-based methane measurements, where repeatability ranged from 0.24 to 0.59 across different measurement intervals and environments (Wahinya et al., 2022). Previously three important environmental factors have been regarded as affecting methane estimates: dietary composition changes (Bica et al., 2022), host physiological variation (Connor et al., 2024), and temporal patterns changes in microbiome diversity (Fenn et al., 2023). For reliable microbiome-methane prediction accuracy, validation strategies should consider the temporal and environmental factors. Leave-one-day-out validation provided a more realistic assessment of prediction accuracy in this study.

### Limitation of methane prediction models

This study provides an evaluation of three microbial feature profiling approaches (SqueezeMeta, Kraken2-NT and Kraken2-GTDB) using BLUP models for methane prediction, however several aspects of the analysis pipeline have not yet been explored. For example, as the majority of previous metagenomic prediction studies have been performed using short read data where the read length is fixed, the effect of the interaction between read length, read number and total data volume on prediction accuracy has not yet been characterized. Likewise, the statistical approaches used have been compared in some cases (Hess et al., 2023; Wang et al., 2024), but not in long read data and were not systematically evaluated. Hence, while this study has characterized the effect of some bioinformatics pipelines on the prediction accuracy of enteric methane in sheep and indeed demonstrated the potential higher prediction accuracy achievable with long-read sequence data, the optimizations of taxonomic and functional feature identification still need to be addressed in future work.

## Conclusion

Functional microbial features (KEGG and COG) consistently achieved higher microbiability and prediction accuracy than taxonomic features across all methane prediction models and validation strategies, demonstrating superior predictive capacity of microbial functional features for methane phenotypes. Different bioinformatic pipelines for taxonomic characterization substantially influenced the prediction model performance. Multi-matrix models integrating both taxonomic and functional features provided only marginal or no improvement in prediction accuracy over single-matrix functional models, indicating that functional features alone capture the majority of predictive information for methane phenotype prediction.

## Materials

### Authors’ contributions

YL and EMR, LTN led the study design, data analysis, interpretation, and manuscript writing. CTO performed the long-read metagenomic sequencing and laboratory experiments and edited the manuscript. PF conducted the methane measurements, collected rumen samples, and PF and MA contributed to data interpretation. YL developed bioinformatic workflows, performed taxonomic and functional annotations, developed prediction models, and conducted all statistical analyses. SY contributed to statistical analyses, model validation strategies, and edited the manuscript. JvdW provided critical datasets and computational resources, co-supervised the project, contributed to interpretation, and edited the manuscript. EMR and LTN secured funding, managed the project including the development of analytical protocols, supervised the research, contributed to interpretation, and edited the manuscript. All authors have read and approved the final manuscript.

### Funding

This research is funded by Meat and Livestock Australia and MLA Donor Company (P.PSH.2010 and P.PSH.2011), the Department of Primary Industries and The University of Queensland. The author acknowledges support through the Australian Government and The University of Queensland Research Training Program (RTP) Scholarship.

## Acknowledgements

The author gratefully acknowledges Collins Asiamah for essential DNA extraction work and thanks Daniel Brown and Ed Clayton for their contribution to this project scope.

