## Additional file 1 for "Metagenomic prediction of methane emissions in sheep using single- and multi-matrix BLUP models with taxonomic and functional microbial features"

**Table S1**. **Principal coordinate analysis (PCoA) and methane production correlation analysis**

| Pipeline | Microbial feature | Principal coordinate | Variance (%) | *p*-value | Significance |
|---|---|---|---|---|---|
| SqueezeMeta | Genus | PCoA1 | 30.79 | 0.2314 | ns |
| | | PCoA2 | 10.36 | 0.1498 | ns |
| | | PCoA3 | 6.79 | 0.4504 | ns |
| | | PCoA4 | 6.27 | 0.5026 | ns |
| | | PCoA5 | 4.57 | 0.0895 | ns |
| Kraken2-NT | Genus | PCoA1 | 56.76 | 0.6394 | ns |
| | | PCoA2 | 6.64 | 0.714 | ns |
| | | PCoA3 | 4.99 | 0.0372 | ns |
| | | PCoA4 | 3.42 | 0.1006 | ns |
| | | PCoA5 | 2.55 | 0.4984 | ns |
| Kraken2-GTDB | Genus | PCoA1 | 37.78 | 0.4674 | ns |
| | | PCoA2 | 8.54 | 0.2616 | ns |
| | | PCoA3 | 4.85 | 0.6773 | ns |
| | | PCoA4 | 4.22 | 0.6937 | ns |
| | | PCoA5 | 3.75 | 0.2381 | ns |
| SqueezeMeta | KEGG | PCoA1 | 72.82 | 0.4518 | ns |
| | | PCoA2 | 4.48 | 0.5659 | ns |
| | | PCoA3 | 2.7 | 0.2935 | ns |
| | | PCoA4 | 2.27 | 0.5253 | ns |
| | | PCoA5 | 1.28 | 0.869 | ns |
| SqueezeMeta | COG | PCoA1 | 59.18 | 0.4287 | ns |
| | | PCoA2 | 6.2 | 0.6196 | ns |
| | | PCoA3 | 4.95 | 0.1202 | ns |
| | | PCoA4 | 2.93 | 0.8126 | ns |
| | | PCoA5 | 2.26 | 0.9346 | ns |

Principal coordinate analysis (PCoA) was performed on genus-, KEGG- and COG-level microbial abundance matrices using Bray–Curtis dissimilarity. Methane production residual values were calculated on body weight and run using linear regression. The first five principal coordinates (PCoA1–5) were analysed with methane production residuals using Spearman rank correlation. Variance (%) indicates the percentage of total variance explained by each principal coordinate. Significance indicates the p-value: ***$p < 0.001$; **$p < 0.01$; *$p < 0.05$; ns, not significant.

**Table S2. Variance components and microbiability estimates in single-matrix BLUP Model**

| Pipeline | Microbial feature | $V_m$ | $V_e$ | Microbiability |
|---|---|---|---|---|
| SqueezeMeta | Phylum | 0.131±0.037 | 0.348±0.031 | 0.273±0.068 |
| | Class | 0.224±0.052 | 0.276±0.033 | 0.448±0.086 |
| | Order | 0.247±0.055 | 0.244±0.036 | 0.503±0.093 |
| | Family | 0.261±0.056 | 0.214±0.038 | 0.549±0.098 |
| | Genus | 0.293±0.058 | 0.168±0.039 | 0.636±0.104 |
| | KEGG | 0.477±0.099 | 0.116±0.043 | 0.805±0.094 |
| | COG | 0.486±0.084 | 0.034±0.045 | 0.935±0.092 |
| Kraken2-NT | Phylum | 0.118±0.039 | 0.349±0.029 | 0.253±0.072 |
| | Class | 0.160±0.049 | 0.325±0.029 | 0.330±0.082 |
| | Order | 0.211±0.056 | 0.284±0.032 | 0.426±0.090 |
| | Family | 0.230±0.065 | 0.263±0.038 | 0.467±0.109 |
| | Genus | 0.342±0.080 | 0.174±0.046 | 0.662±0.116 |
| Kraken2-GTDB | Phylum | 0.255±0.065 | 0.331±0.029 | 0.435±0.077 |
| | Class | 0.275±0.071 | 0.301±0.034 | 0.478±0.091 |
| | Order | 0.358±0.089 | 0.228±0.046 | 0.611±0.112 |
| | Family | 0.393±0.100 | 0.190±0.057 | 0.674±0.129 |
| | Genus | 0.439±0.097 | 0.116±0.061 | 0.791±0.131 |

All variance estimates are presented as mean ± standard error from BLUP models. $V_m$ ($\sigma_m^2$) represents the microbial feature variance component. $V_e$ ($\sigma_e^2$) represents the residual variance. Microbiability ($m^2$) represents the proportion of phenotypic variance explained by rumen microbiome composition, calculated as $m^2 = \sigma_m^2 / (\sigma_m^2 + \sigma_e^2)$. Higher microbiability values indicate stronger associations between microbial features and methane emissions.

**Table S3. Variance components and microbiability estimates multi-matrices BLUP Model**

| Pipeline | Microbial features | $V_t$ (Taxon) | $V_f$ (Function) | $V_e$ | Microbiability |
|---|---|---|---|---|---|
| SqueezeMeta | Phylum + KEGG | 0.000±0.000 | 0.477±0.103 | 0.116±0.043 | 0.805±0.094 |
| | Class + KEGG | 0.020±0.037 | 0.454±0.105 | 0.113±0.043 | 0.808±0.094 |
| | Order + KEGG | 0.019±0.043 | 0.456±0.108 | 0.112±0.043 | 0.809±0.094 |
| | Family + KEGG | 0.019±0.049 | 0.456±0.111 | 0.111±0.044 | 0.810±0.095 |
| | Genus + KEGG | 0.071±0.057 | 0.404±0.108 | 0.095±0.044 | 0.833±0.094 |
| Kraken2-NT, SqueezeMeta (KEGG) | Phylum + KEGG | 0.071±0.057 | 0.404±0.108 | 0.095±0.044 | 0.833±0.094 |
| | Class + KEGG | 0.034±0.027 | 0.427±0.100 | 0.121±0.044 | 0.793±0.099 |
| | Order + KEGG | 0.027±0.030 | 0.419±0.102 | 0.128±0.044 | 0.777±0.101 |
| | Family + KEGG | 0.078±0.045 | 0.386±0.099 | 0.112±0.043 | 0.805±0.096 |
| | Genus + KEGG | 0.058±0.050 | 0.400±0.104 | 0.116±0.044 | 0.798±0.099 |
| Kraken2-GTDB, SqueezeMeta (KEGG) | Phylum + KEGG | 0.069±0.063 | 0.388±0.108 | 0.113±0.045 | 0.802±0.100 |
| | Class + KEGG | 0.037±0.037 | 0.450±0.098 | 0.111±0.042 | 0.814±0.092 |
| | Order + KEGG | 0.002±0.028 | 0.475±0.101 | 0.115±0.043 | 0.805±0.095 |
| | Family + KEGG | 0.000±0.000 | 0.477±0.102 | 0.116±0.044 | 0.805±0.095 |
| | Genus + KEGG | 0.000±0.000 | 0.477±0.099 | 0.116±0.043 | 0.805±0.094 |
| SqueezeMeta | Phylum + COG | 0.000±0.000 | 0.487±0.084 | 0.034±0.045 | 0.935±0.092 |
| | Class + COG | 0.027±0.034 | 0.465±0.087 | 0.027±0.045 | 0.947±0.092 |
| | Order + COG | 0.032±0.039 | 0.464±0.088 | 0.024±0.045 | 0.954±0.092 |
| | Family + COG | 0.045±0.045 | 0.451±0.089 | 0.021±0.046 | 0.960±0.092 |
| | Genus + COG | 0.080±0.050 | 0.430±0.071 | 0.013±0.045 | 0.975±0.086 |
| Kraken2-NT, SqueezeMeta (COG) | Phylum + COG | 0.038±0.028 | 0.447±0.080 | 0.040±0.046 | 0.924±0.094 |
| | Class + COG | 0.031±0.029 | 0.446±0.082 | 0.044±0.046 | 0.916±0.096 |
| | Order + COG | 0.094±0.044 | 0.432±0.058 | 0.027±0.045 | 0.951±0.082 |
| | Family + COG | 0.067±0.049 | 0.438±0.073 | 0.032±0.046 | 0.940±0.089 |
| | Genus + COG | 0.068±0.059 | 0.428±0.082 | 0.034±0.046 | 0.936±0.092 |
| Kraken2-GTDB, SqueezeMeta (COG) | Phylum + COG | 0.050±0.038 | 0.466±0.070 | 0.031±0.044 | 0.944±0.085 |
| | Class + COG | 0.015±0.038 | 0.474±0.086 | 0.033±0.045 | 0.937±0.092 |
| | Order + COG | 0.001±0.025 | 0.496±0.074 | 0.034±0.045 | 0.936±0.089 |
| | Family + COG | 0.000±0.000 | 0.498±0.070 | 0.034±0.045 | 0.936±0.089 |
| | Genus + COG | 0.000±0.000 | 0.486±0.084 | 0.034±0.045 | 0.935±0.092 |

A multi-matrix BLUP model was used to simultaneously estimate variance components for methane emissions and the associated microbial features. The model partitions the total variance into: $V_t$ ($\sigma_t^2$), the variance attributable to microbial taxonomic classification; $V_f$ ($\sigma_f^2$), the variance attributable to functional annotations; and $V_e$ ($\sigma_e^2$), the residual variance. Microbiability ($m^2$) represents the proportion of phenotypic variance explained by the microbial feature relationship matrices. All estimates are presented as mean ± standard error.

**Table S4. 5-fold cross-validation results**

| Pipeline | Microbial features | Accuracy (*r*) | RMSE |
|---|---|---|---|
| SqueezeMeta | Phylum | 0.444±0.021 | 0.917±0.026 |
| | Class | 0.498±0.025 | 0.896±0.027 |
| | Order | 0.545±0.010 | 0.879±0.027 |
| | Family | 0.545±0.019 | 0.879±0.026 |
| | Genus | 0.575±0.015 | 0.860±0.025 |
| | KEGG | 0.624±0.032 | 0.815±0.026 |
| | COG | 0.634±0.038 | 0.806±0.031 |
| Kraken2-NT | Phylum | 0.436±0.054 | 0.915±0.020 |
| | Class | 0.481±0.042 | 0.893±0.018 |
| | Order | 0.487±0.047 | 0.893±0.017 |
| | Family | 0.514±0.041 | 0.884±0.020 |
| | Genus | 0.529±0.045 | 0.875±0.021 |
| Kraken2-GTDB | Phylum | 0.429±0.052 | 0.917±0.023 |
| | Class | 0.413±0.049 | 0.928±0.023 |
| | Order | 0.487±0.049 | 0.911±0.022 |
| | Family | 0.502±0.040 | 0.912±0.022 |
| | Genus | 0.533±0.031 | 0.906±0.022 |
| SqueezeMeta | Phylum + KEGG | 0.621±0.033 | 0.817±0.026 |
| | Class + KEGG | 0.624±0.031 | 0.814±0.025 |
| | Order + KEGG | 0.626±0.031 | 0.813±0.025 |
| | Family + KEGG | 0.625±0.031 | 0.814±0.025 |
| | Genus + KEGG | 0.629±0.029 | 0.812±0.025 |
| Kraken2-NT, SqueezeMeta (KEGG) | Phylum + KEGG | 0.604±0.035 | 0.823±0.025 |
| | Class + KEGG | 0.613±0.030 | 0.820±0.023 |
| | Order + KEGG | 0.622±0.030 | 0.815±0.022 |
| | Family + KEGG | 0.624±0.032 | 0.815±0.025 |
| | Genus + KEGG | 0.619±0.030 | 0.817±0.024 |
| Kraken2-GTDB, SqueezeMeta (KEGG) | Phylum + KEGG | 0.618±0.033 | 0.817±0.026 |
| | Class + KEGG | 0.621±0.031 | 0.817±0.025 |
| | Order + KEGG | 0.623±0.032 | 0.816±0.025 |
| | Family + KEGG | 0.624±0.032 | 0.815±0.026 |
| | Genus + KEGG | 0.623±0.032 | 0.816±0.025 |
| SqueezeMeta | Phylum + COG | 0.632±0.039 | 0.807±0.032 |
| | Class + COG | 0.636±0.037 | 0.804±0.031 |
| | Order + COG | 0.641±0.035 | 0.801±0.031 |
| | Family + COG | 0.640±0.035 | 0.803±0.031 |
| | Genus + COG | 0.645±0.034 | 0.800±0.030 |
| Kraken2-NT, | Phylum + COG | 0.616±0.039 | 0.814±0.029 |

| | | | |
|---|---|---|---|
| SqueezeMeta (COG) | Class + COG | 0.627±0.035 | 0.809±0.027 |
| | Order + COG | 0.637±0.033 | 0.803±0.025 |
| | Family + COG | 0.638±0.036 | 0.804±0.029 |
| | Genus + COG | 0.634±0.035 | 0.806±0.029 |
| Kraken2-GTDB, SqueezeMeta (COG) | Phylum + COG | 0.627±0.040 | 0.809±0.031 |
| | Class + COG | 0.633±0.037 | 0.806±0.031 |
| | Order + COG | 0.634±0.037 | 0.806±0.031 |
| | Family + COG | 0.634±0.038 | 0.806±0.031 |
| | Genus + COG | 0.634±0.038 | 0.806±0.031 |

Mean prediction accuracy (Pearson correlation coefficient, *r*) and root mean square error (RMSE) for single-matrix and multi-matrix BLUP models using different bioinformatic pipelines for generating microbial features. Prediction accuracy (*r*) and RMSE, presented as mean ± standard error across five cross-validation folds, are shown for taxonomic levels (phylum to genus) and functional annotations (KEGG and COG) from the SqueezeMeta, Kraken2-NT, and Kraken2-GTDB pipelines.

**Table S5. Leave-one-day-out validation results**

| Pipeline | Microbial features | Accuracy (*r*) | RMSE |
| --- | --- | --- | --- |
| SqueezeMeta | Phylum | 0.360±0.042 | 0.959±0.042 |
| | Class | 0.452±0.040 | 0.935±0.043 |
| | Order | 0.468±0.042 | 0.926±0.039 |
| | Family | 0.451±0.043 | 0.931±0.037 |
| | Genus | 0.474±0.047 | 0.915±0.034 |
| | KEGG | 0.535±0.037 | 0.866±0.033 |
| | COG | 0.544±0.035 | 0.851±0.034 |
| Kraken2-NT | Phylum | 0.352±0.035 | 0.940±0.039 |
| | Class | 0.416±0.030 | 0.927±0.037 |
| | Order | 0.399±0.027 | 0.935±0.043 |
| | Family | 0.413±0.032 | 0.922±0.037 |
| | Genus | 0.414±0.031 | 0.917±0.037 |
| Kraken2-GTDB | Phylum | 0.329±0.048 | 0.963±0.036 |
| | Class | 0.322±0.050 | 0.969±0.039 |
| | Order | 0.379±0.034 | 0.950±0.040 |
| | Family | 0.372±0.049 | 0.954±0.039 |
| | Genus | 0.428±0.039 | 0.945±0.037 |
| SqueezeMeta | Phylum + KEGG | 0.535±0.037 | 0.867±0.033 |
| | Class + KEGG | 0.539±0.038 | 0.866±0.032 |
| | Order + KEGG | 0.536±0.038 | 0.867±0.032 |
| | Family + KEGG | 0.536±0.038 | 0.867±0.033 |
| | Genus + KEGG | 0.545±0.039 | 0.864±0.032 |
| Kraken2-NT, SqueezeMeta (KEGG) | Phylum + KEGG | 0.526±0.037 | 0.867±0.031 |
| | Class + KEGG | 0.539±0.036 | 0.866±0.031 |
| | Order + KEGG | 0.554±0.031 | 0.861±0.034 |
| | Family + KEGG | 0.542±0.036 | 0.864±0.031 |
| | Genus + KEGG | 0.537±0.033 | 0.865±0.033 |
| Kraken2-GTDB, SqueezeMeta (KEGG) | Phylum + KEGG | 0.532±0.036 | 0.869±0.032 |
| | Class + KEGG | 0.533±0.036 | 0.869±0.033 |
| | Order + KEGG | 0.534±0.036 | 0.867±0.033 |
| | Family + KEGG | 0.535±0.037 | 0.867±0.033 |
| | Genus + KEGG | 0.535±0.037 | 0.867±0.033 |
| SqueezeMeta | Phylum + COG | 0.551±0.040 | 0.870±0.034 |
| | Class + COG | 0.547±0.035 | 0.851±0.034 |
| | Order + COG | 0.541±0.038 | 0.853±0.032 |
| | Family + COG | 0.544±0.037 | 0.853±0.033 |
| | Genus + COG | 0.553±0.037 | 0.849±0.033 |
| Kraken2-NT. SqueezeMeta (COG) | Phylum + COG | 0.549±0.034 | 0.867±0.035 |
| | Class + COG | 0.545±0.035 | 0.851±0.033 |

| | | | |
|---|---|---|---|
| | Order + COG | 0.559±0.033 | 0.866±0.035 |
| | Family + COG | 0.558±0.039 | 0.864±0.033 |
| | Genus + COG | 0.543±0.033 | 0.852±0.034 |
| Kraken2-GTDB, SqueezeMeta (COG) | Phylum + COG | 0.537±0.033 | 0.854±0.033 |
| | Class + COG | 0.542±0.033 | 0.853±0.034 |
| | Order + COG | 0.549±0.039 | 0.871±0.035 |
| | Family + COG | 0.551±0.040 | 0.870±0.034 |
| | Genus + COG | 0.551±0.040 | 0.871±0.034 |

Mean prediction accuracy (Pearson correlation coefficient, *r*) and root mean square error (RMSE) for single-matrix and multi-matrix BLUP models using leave-one-day-out cross-validation. Each measurement day served sequentially as the test set while the remaining six days formed the training set. Prediction accuracy (*r*) and RMSE are presented as mean ± standard error across seven validation folds (one for each day).
