## Additional file 2 for "Metagenomic prediction of methane emissions in sheep using single- and multi-matrix BLUP models with taxonomic and functional microbial features"

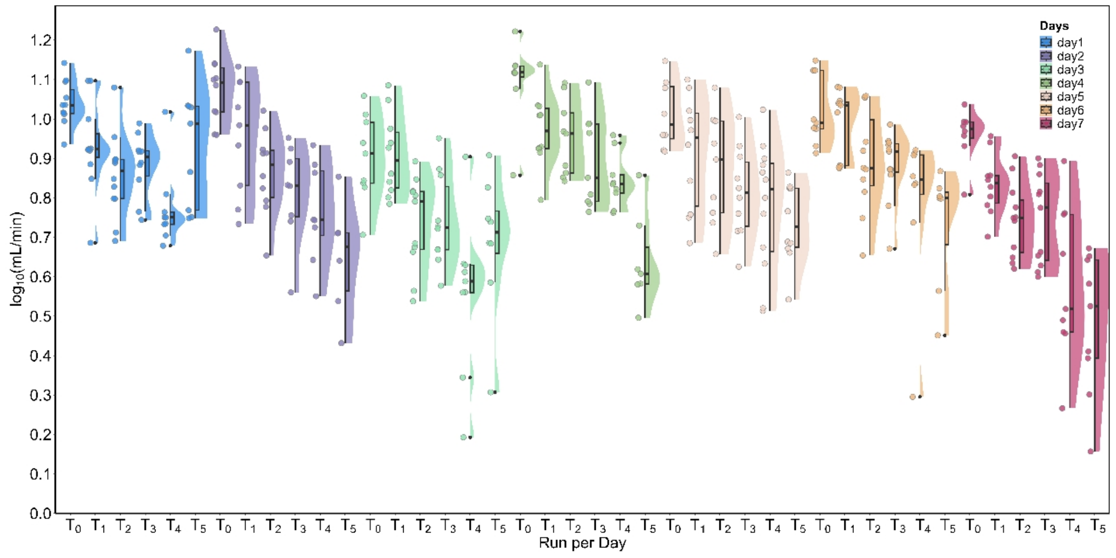


**Fig. S1 Methane production (mL/min) distribution across six sampling time points and seven days.**

Methane production showed a clear temporal pattern across runs. The raincloud plot displays methane production across 6 runs and 7 days (X-axis). Each visualization combines half-violin plots representing methane density distributions, with box plots indicating median values with interquartile ranges and individual points representing $\log_{10}$ transformed methane production (Y-axis). Each run represents specific methane measurement time: $T_0$ (08:00–08:16), $T_1$ (09:30–09:46), $T_2$ (11:00–11:16), $T_3$ (12:30–12:46), $T_4$ (14:00–14:16), and $T_5$ (15:30–15:46), which correspond to different colors representing measurement days.

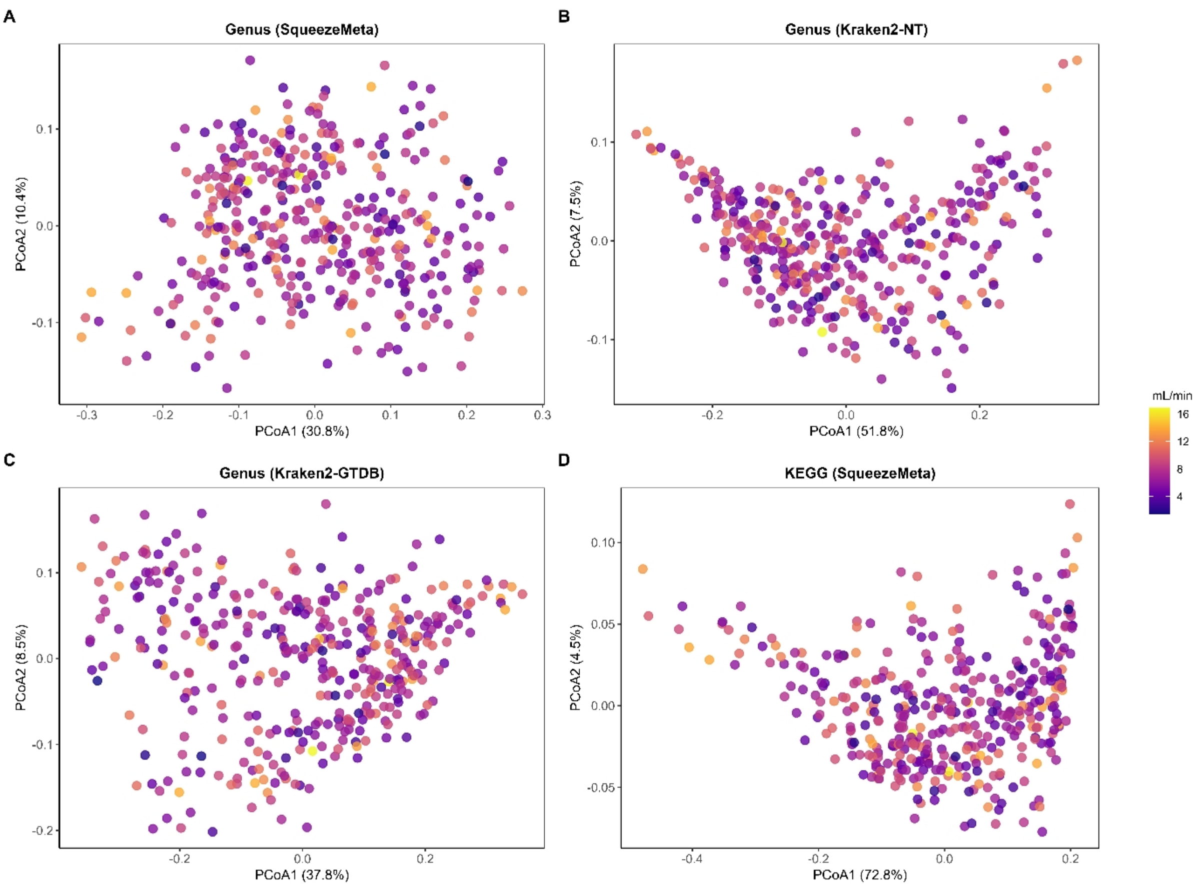


**Fig S2. Principal coordinate analysis (PCoA) of rumen microbiome feature diversity associated with methane production phenotypes.**

(A) Genus-level taxonomic abundance profiled using SqueezeMeta, (B) Genus-level taxonomic abundance profiled using Kraken2-NT, (C) Genus-level taxonomic abundance profiled using Kraken2-GTDB, (D) KEGG functional abundance profiled using SqueezeMeta. Each point represents one animal. Points are colored by methane production (mL/min).
