## Additional file 3 for "Metagenomic prediction of methane emissions in sheep using single- and multi-matrix BLUP models with taxonomic and functional microbial features"

**Additional file 3.** Feature-wise shuffling of microbial abundance matrices as a negative control.

### Supplementary Methods

To test for an unknown effect of the data structure on microbiability or prediction accuracy, the count matrices were randomized to provide a null control. Because randomization of the count matrix should remove the relationship between the microbiome data and the phenotype, we hypothesized that microbiability and prediction accuracy would not differ significantly from zero when the randomized matrix was used. To test this hypothesis, random shuffling was applied feature-wise to all microbial matrices as a negative control. Within each taxonomic or functional feature, the abundance values were randomly reassigned across all animals, so that every animal received a randomly assigned value drawn from the original distribution of that feature. This produced, for each count matrix, a new matrix of randomly shuffled values, which was then used to construct a microbial relationship matrix. The relationship matrices generated from the shuffled data were used to estimate microbiability and prediction accuracy (under both 5-fold and leave-one-day-out cross-validation), applying identical model specifications to those used for the non-shuffled data.

#### Supplementary Results

Feature-wise random shuffling of the microbial matrices collapsed both microbiability and prediction accuracy to values that did not differ significantly from zero across all BLUP models (Table S6). The shuffled matrices broke the association between microbial composition and the methane phenotype. These results confirmed that the microbial signal in the original analyses reflects a genuine link between the count information and the phenotype, rather than an artefact of matrix size or uneven coverage of features.

**Table S6.** Microbiability ($m^2$) and mean prediction accuracy (*r*, 5-fold cross-validation and leave-one-day-out validation) estimated from microbial relationship matrices constructed after feature-wise random shuffling. For single-matrix models, the taxonomic or functional matrix was shuffled; for multi-matrix models, both matrices were shuffled. Values are reported as mean ± standard error.

| Model | Pipeline | Microbial Feature | Microbiability ($m^2$) | Accuracy (*r*) | |
|---|---|---|---|---|---|
| | | | | 5-fold CV | Leave-one-day-out |
| Single-matrix | SqueezeMeta | Phylum | 0.000±0.053 | 0.022±0.037 | 0.025±0.048 |
| | | Class | 0.028±0.092 | -0.025±0.065 | 0.065±0.043 |
| | | Order | 0.021±0.114 | -0.060±0.053 | -0.074±0.050 |

| | | | | | |
|---|---|---|---|---|---|
| | | Family | 0.121±0.142 | -0.000±0.042 | 0.037±0.056 |
| | | Genus | 0.116±0.184 | -0.011±0.042 | -0.020±0.072 |
| | | KEGG | 0.000±0.360 | 0.010±0.030 | -0.048±0.028 |
| | | COG | 0.000±0.540 | 0.033±0.042 | 0.059±0.057 |
| | Kraken2-NT | Phylum | 0.053±0.063 | 0.016±0.053 | 0.161±0.042 |
| | | Class | 0.000±0.034 | 0.014±0.035 | -0.067±0.058 |
| | | Order | 0.000±0.071 | -0.056±0.016 | -0.121±0.066 |
| | | Family | 0.101±0.111 | 0.075±0.041 | 0.046±0.051 |
| | | Genus | 0.162±0.188 | -0.063±0.057 | 0.105±0.040 |
| | Kraken2-GTDB | Phylum | 0.000±0.050 | 0.000±0.030 | 0.000±0.046 |
| | | Class | 0.037±0.092 | -0.030±0.055 | -0.043±0.014 |
| | | Order | 0.000±0.145 | -0.071±0.034 | -0.033±0.040 |
| | | Family | 0.000±0.235 | -0.088±0.065 | -0.105±0.038 |
| | | Genus | 0.178±0.306 | 0.018±0.060 | 0.014±0.045 |
| Multi-matrix | SqueezeMeta | Phylum + KEGG | 0.000±0.053 | 0.010±0.030 | -0.022±0.037 |
| | | Class + KEGG | 0.028±0.373 | 0.196±0.042 | 0.166±0.034 |
| | | Order + KEGG | 0.021±0.114 | 0.007±0.053 | -0.015±0.042 |
| | | Family + KEGG | 0.121±0.407 | -0.010±0.039 | -0.105±0.046 |
| | | Genus + KEGG | 0.116±0.421 | -0.011±0.041 | -0.052±0.034 |
| | Kraken2-NT + SqueezeMeta (KEGG) | Phylum + KEGG | 0.052±0.063 | -0.007±0.035 | -0.004±0.048 |
| | | Class + KEGG | 0.000±0.034 | -0.073±0.035 | -0.090±0.037 |
| | | Order + KEGG | 0.000±0.000 | -0.012±0.025 | -0.082±0.038 |
| | | Family + KEGG | 0.101±0.404 | 0.025±0.094 | 0.027±0.052 |
| | | Genus + KEGG | 0.162±0.188 | 0.057±0.040 | 0.055±0.042 |
| | Kraken2-GTDB + SqueezeMeta (KEGG) | Phylum + KEGG | 0.000±0.050 | 0.026±0.022 | -0.035±0.046 |
| | | Class + KEGG | 0.036±0.091 | -0.032±0.044 | -0.048±0.015 |
| | | Order + KEGG | 0.000±0.000 | -0.050±0.014 | -0.035±0.034 |
| | | Family + KEGG | 0.000±0.000 | -0.021±0.064 | -0.074±0.029 |
| | | Genus + KEGG | 0.178±0.477 | -0.011±0.053 | 0.001±0.041 |
| Multi-matrix | SqueezeMeta | Phylum + COG | 0.000±0.000 | -0.010±0.043 | -0.022±0.044 |
| | | Class + COG | 0.279±0.104 | 0.197±0.041 | 0.164±0.039 |
| | | Order + COG | 0.062±0.557 | -0.010±0.045 | -0.037±0.053 |
| | | Family + COG | 0.000±0.000 | -0.041±0.031 | -0.080±0.049 |
| | | Genus + COG | 0.000±0.169 | -0.047±0.028 | -0.056±0.035 |
| | Kraken2-NT + SqueezeMeta (COG) | Phylum + COG | 0.203±0.565 | 0.068±0.042 | 0.090±0.052 |
| | | Class + COG | 0.201±0.560 | -0.022±0.066 | -0.015±0.069 |
| | | Order + COG | 0.193±0.558 | 0.003±0.041 | 0.013±0.065 |
| | | Family + COG | 0.193±0.565 | 0.025±0.092 | 0.034±0.043 |
| | | Genus + COG | 0.283±0.579 | 0.051±0.039 | 0.073±0.049 |
| | Kraken2-GTDB + SqueezeMeta (COG) | Phylum + COG | 0.000±0.050 | 0.017±0.038 | 0.056±0.067 |
| | | Class + COG | 0.036±0.091 | -0.049±0.029 | 0.007±0.034 |
| | | Order + COG | 0.000±0.000 | -0.008±0.053 | 0.016±0.042 |
| | | Family + COG | 0.000±0.000 | -0.036±0.066 | -0.056±0.042 |

| | | | | |
|---|---|---|---|---|
| | Genus + COG | 0.178±0.649 | 0.016±0.061 | -0.013±0.041 |
